# Identification of Dehydrogenase, Hydratase, and Aldolase Required for C17 Propionyl Residue Removal in Steroid Degradation by *Comamonas testosteroni* TA441: Structural and Functional Insights into the (ChsE1-ChsE2)_2_ and (ChsH1-ChsH2_MaoC_)_2_-(Ltp2-ChsH2_DUF35_)_2_ Complexes

**DOI:** 10.1101/2025.01.07.631791

**Authors:** Masae Horinouchi

**Author notes:** Address correspondence to Masae Horinouchi.

## Abstract

Bacterial steroid degradation is gaining attention for its diverse roles, such as *Mycobacterium tuberculosis*’s reliance on the degradation of the C17 side chain of cholesterol for survival in host environments. ORF40 to ORF44 of *Comamonas testosteroni* TA441, previously characterized for its A-, B-, C-, and D-ring cleavage pathways, were hypothesized to correspond to the *igr* operon of *M. tuberculosis*, encoding the dehydrogenase ChsE1E2, hydratase ChsH1H2 _maoC_, and aldolase Ltp2 ChsH2_DUF35_, responsible for propionyl residue removal in C17 side-chain degradation. However, low amino acid identity between the corresponding enzymes precluded functional assignment without experimental evidence.

In this study, we generated gene-disrupted mutants of ORF40–44 and demonstrated that ORF41/44, ORF40/42, and ORF43 encode the dehydrogenase, hydratase, and aldolase, respectively. ORF40 encodes a bifunctional protein comprising MaoC and DUF35 domains. The MaoC domain of ORF40 and the ORF42-encoded protein form the hydratase, while the DUF35 domain is essential for aldolase activity. A mutant expressing ORF40_MaoC_ and ORF40_DUF35_ separately exhibited both hydratase and aldolase activities, suggesting these activities do not require a strictly formed complex of hydratase and aldolase. However, efficient propionyl residue removal appears to depend on the proper formation of each enzymatic complex, including ChsE1E2, ChsH1H2_MaoC_, and Ltp2ChsH2_DUF35_; although (ChsE1-ChsE2)_2_ does not form a stable complex with (ChsH1-ChsH2_MaoC_)_2_-(Ltp2-ChsH2_DUF35_)_2_, some degree of interaction was suggested. AlphaFold-predicted 3D structures of the TA441 enzymes and their complexes revealed striking similarities to those of *M. tuberculosis*, despite low amino acid identities. These findings shed light on the structural and functional conservation of bacterial steroid-degrading enzymes.

**IMPORTANCE:** Research on bacterial steroid degradation began over 50 years ago, primarily to produce substrates for steroid drugs. Recently, the role of steroid-degrading bacteria in human health has garnered increasing attention. *Comamonas testosteroni* TA441 is a prominent model organism for studying aerobic steroid degradation, with its overall pathways for A-, B-, C-, and D-ring cleavage already elucidated. In this study, we identified the mechanism for removing the propionyl residue at C17, a crucial step in degrading steroids with a C17 side chain, such as cholic acid, cholesterol, and other biologically significant compounds in animals and plants. The functions and structures of the identified enzymes show remarkable similarity to those in *Mycobacterium tuberculosis*. These findings suggest that insights gained from TA441 could provide valuable clues for understanding *M. tuberculosis* steroid metabolism and the broader ecological and health-related implications of bacterial steroid degradation.

## INTRODUCTION

Steroid compounds perform various functions in both plants and animals, including humans, where they play critical roles such as hormone, cholesterol, and bile acid. These compounds have garnered increasing attention due to their impact on human health, particularly in the context of pathogenic bacteria. For example, a cholesterol import system is essential for the persistence of *Mycobacterium tuberculosis* H37Rv in the lungs of chronically infected animals (1), and cholesterol catabolism is crucial for pathogen maintenance in the host (2).

Cholic acid and deoxycholic acid, the primary components of human bile acids, are secreted into the bile and later reabsorbed in the ileum, where approximately 95% of bile acids are reabsorbed. The remaining portion reaches the colon, where it is converted into secondary bile acids by resident bacteria, influencing human health. Recent studies have revealed the presence of bacterial flora in the jejunum and ileum, suggesting bile acid transformation by bacteria in these regions (3). The bacterial composition in the ileum differs from that in the colon, with ileal microbiotas showing dynamic adaptation to fluctuating intraluminal ecological conditions. However, studies on ileal bacteria are limited due to sampling difficulties.

The ability of bacteria to aerobically degrade steroids has been known for over 70 years, with pioneering studies on *Rhodococcus equi* and *Comamonas testosteroni* conducted in the 1960s (4–8). These studies identified major intermediates in the A- and B-ring degradation pathways and revealed a similar mechanism in both bacteria. Our previous research outlined the degradation pathways for the steroidal four rings (A-, B-, C-, and D-rings) in *C. testosteroni* TA441, encoded within a 120-kb mega-cluster of steroid degradation genes (Fig. 1B). The process involves B-ring cleavage and A-ring aromatization (9–16), followed by A-ring cleavage and D- and C-ring degradation primarily via β-oxidation (17–21). This pathway has established *C. testosteroni* TA441 as a leading model organism for studying bacterial aerobic steroid degradation. Similar pathways are expected in other aerobic steroid-degrading bacteria, including Rhodococci and Mycobacteria.

**Fig. 1.**
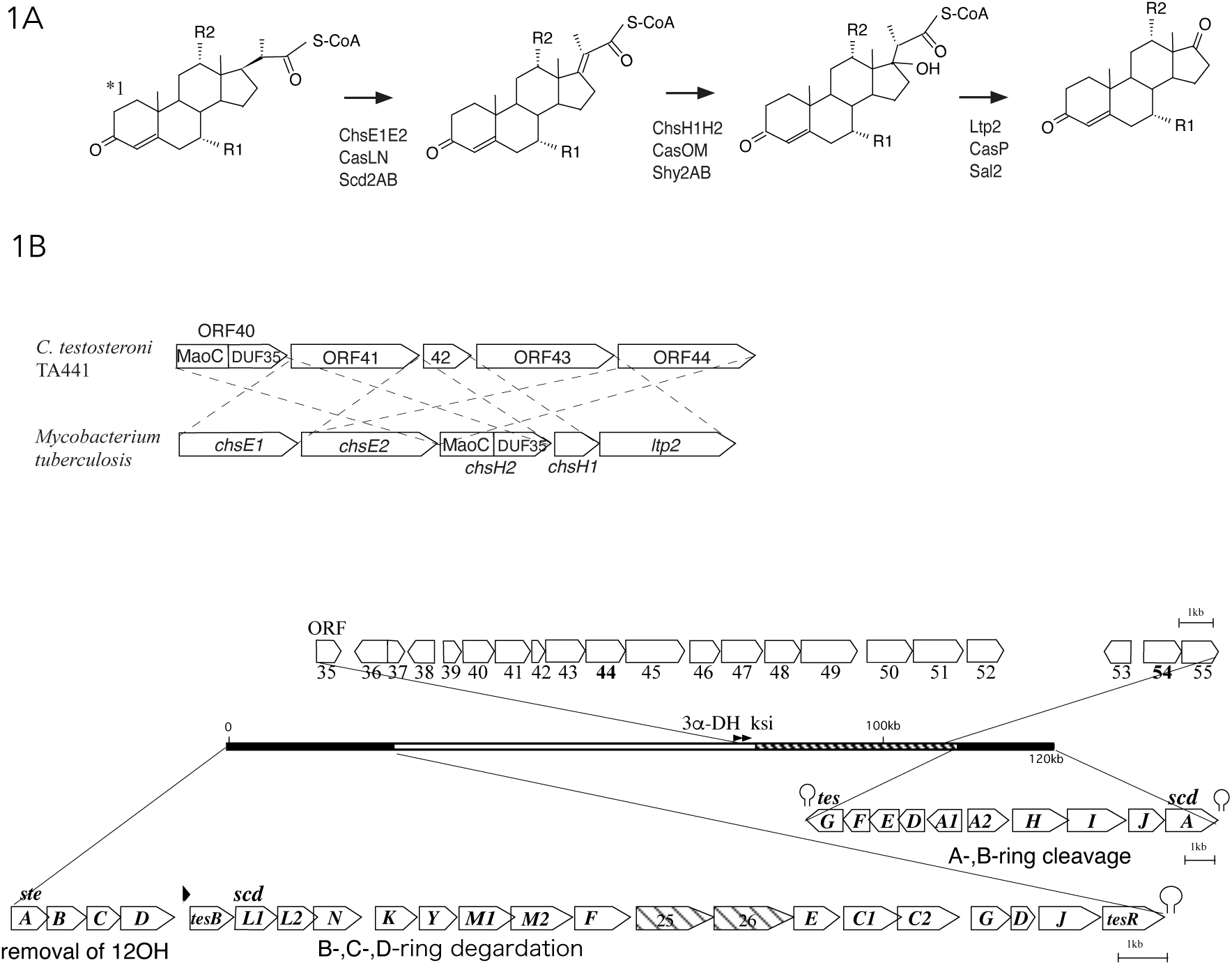
(A) The degradation process of the C20–22 propionyl residue in aerobic steroid degradation reported in Actinobacteria (*Mycobacterium tuberculosis* H37Rv (2, 22–26), *Rhodococcus jostii* RHA1 (27, 28), *Thermomonospora curvata* (29), and *Mycolicibacterium neoaurum* DSM 44074 30)) and the Proteobacterium *Pseudomonas stutzeri* Chol1 (31). The initial reaction is catalyzed by a dehydrogenase (ChsE1E2 in *M. tuberculosis*, *T. curvata*, and *M. neoaurum*; CasLN in *R. jostii* RHA1; and Scd2AB in *P. stutzeri*), followed by the addition of a water molecule by a hydratase (ChsH1H2 in *M. tuberculosis*, *T. curvata*, and *M. neoaurum*; CasOM in *R. jostii* RHA1; and Shy2AB in *P. stutzeri*). Finally, the propionyl residue is removed by an aldolase (Ltp2 in *M. tuberculosis*, *T. curvata*, and *M. neoaurum*; CasP in *R. jostii* RHA1; and Sal2 in *P. stutzeri*). *1: The A-ring has a double bond at C4 in Actinobacteria and two double bonds at C1 and C4 in *P. stutzeri*. R1 and R2 represent H or OH groups. (B) Comparison of *chsE1*, *E2*, *H1*, *H2*, and *ltp2* in *M. tuberculosis* with ORF40 to 44 in *C. testosteroni* TA441 (upper) and the steroid degradation gene mega-cluster of TA441 (bottom).

*C. testosteroni* TA441 degrades various steroids, such as testosterone and cholic acid, with remarkable efficiency. While the pathways for the steroid rings have been characterized, the mechanisms for degrading the C17 side chain remain unclear. In previous studies, we identified intermediate compounds from the culture of a TA441 mutant cultivated with cholic acid (Fig. 2) (17). These intermediates suggest that C17 side-chain degradation occurs concurrently with A-ring dehydrogenation. However, some intermediates were too scarce to identify, and one spontaneously converted to compound 7α,12α-dihydroxy-5β-cholan-4,17-dien-24-oic acid (**VII**) during storage. Cholic acid C17 side-chain degradation involves two main steps: removal of C23 and C24, followed by the removal of the C20-22 propionyl residue. The conversion of the C20-22 propionyl residue to a ketone is a fundamental process shared among bacteria capable of degrading steroids such as cholesterol, cholic acid, and ergosterol. This process has been described in actinobacteria such as *M. tuberculosis* H37Rv (2, 22–26), *Rhodococcus jostii* RHA1 (27, 28), *Thermomonospora curvata* (29), and *Mycolicibacterium neoaurum* DSM 44074 (30), as well as the proteobacterium *Pseudomonas stutzeri* Chol1 (31) (Fig. 1A).

**Fig. 2.**
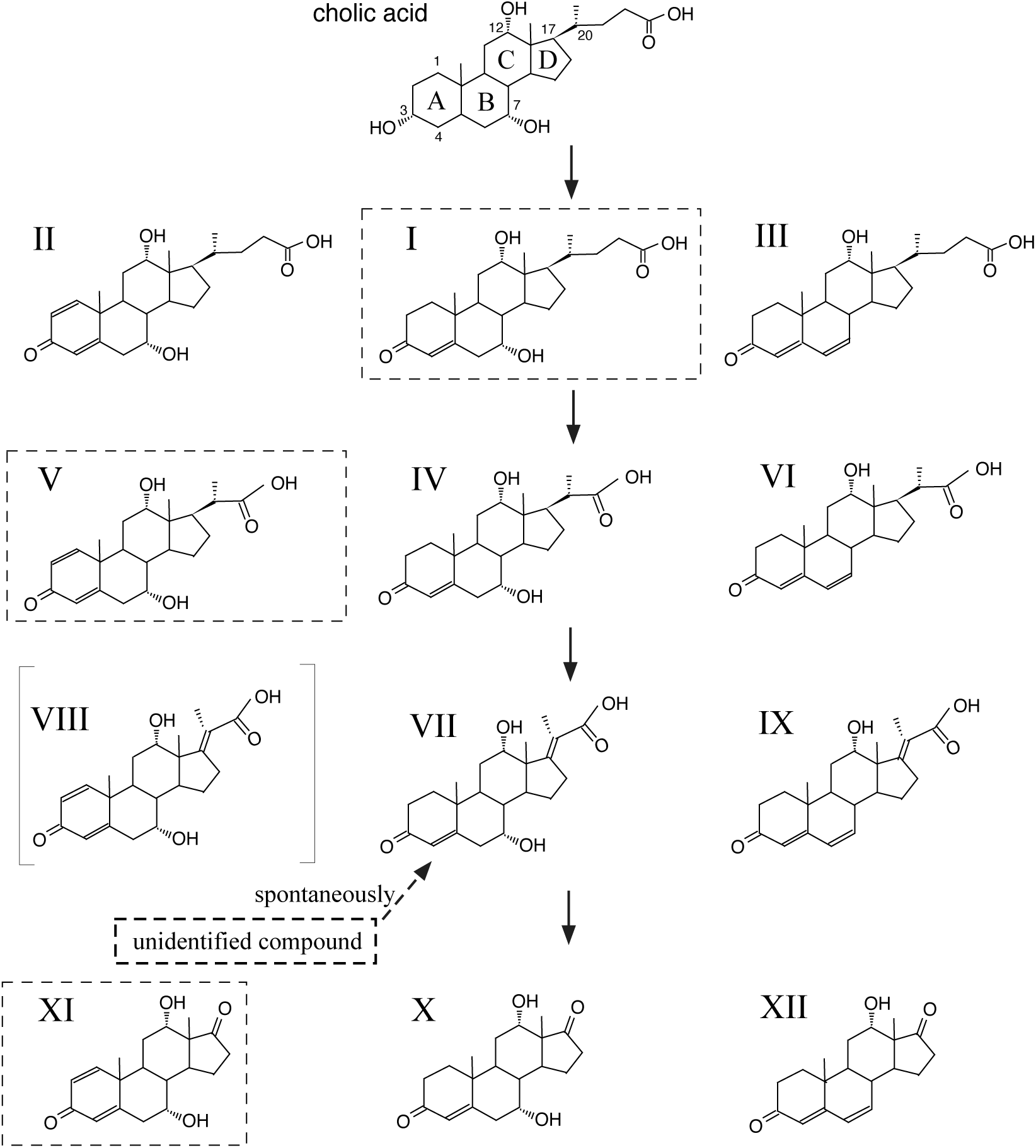
Compounds identified as metabolites of cholic acid during C17 side-chain degradation by *C. testosteroni* TA441 in a previous study. Compounds were isolated and identified using NMR and high-resolution mass spectrometry (17). The actual metabolites in the degradation process are CoA esters. Compounds within broken squares were isolated in larger amounts than others with the same C17 side chain and are considered major metabolites. The compounds are: cholic acid (3α,7α,12α-trihydroxy-5β-cholan-24-oic acid); 7α,12α-dihydroxy-3-oxo-5β-4-cholen-24-oic acid (**I**); 7α,12α-dihydroxy-3-oxo-5β-1,4-choladien-24-oic acid (**II**); 12α-hydroxy-3-oxo-5β-4,6-choladien-24-oic acid (**III**); 7α,12α-dihydroxy-3-oxo-4-pregnene-20-carboxylic acid (**IV**); 7α,12α-dihydroxy-3-oxo-1,4-pregnadine-20-carboxylic acid (**V**); 12α-hydroxy-3-oxo-4,6-pregnadiene-20-carboxylic acid (**VI**); 7α,12α-dihydroxy-3-oxo-4,17-pregnadiene-20-carboxylic acid (**VII**); 7α,12α-dihydroxy-3-oxo-1,4,17-pregnatrine-20-carboxylic acid (**VIII**); 12α-hydroxy-3-oxo-4,6,17-pregnatrine-20-carboxylic acid (**IX**); 7α,12α-dihydroxy-3,17-dioxo-4-androsten (**X**); 7α,12α-dihydroxy-3,17-dioxo-1,4-androstadien (**XI**); and 12α-dihydroxy-3,17-dioxo-4,6-androstadien (**XII**).

The degradation of the C20-22 propionyl residue was first detailed in *M. tuberculosis* H37Rv (22–26). This involves the introduction of a double bond at C17-C20 by a dehydrogenase, followed by hydroxyl group addition at C17 by a hydratase. The isopropionyl residue is then cleaved by an aldolase, producing a ketone at C17 (Fig. 1A). The hydratase in *M. tuberculosis* is a heterotetramer of ChsH1 and MaoC-like protein from the N-terminal domain of ChsH2 (Fig. 1B upper) (25). The aldolase is a heterotetrameric complex of Ltp2 and a DUF35 protein derived from the C-terminal domain of ChsH2 (29).

Based on intermediate compounds isolated from *C. testosteroni* TA441 and observations of a mutant with transposon in ORF44 accumulating substrates of the dehydrogenase (17), we hypothesized that TA441 utilizes a similar pathway. However, the low amino acid identity (30-45%) between TA441’s proteins encoded ORFs near ORF44 and the corresponding *M. tuberculosis* proteins prevented functional assignment without experimental evidence. Here, we report the identification and characterization of the C20-22 propionyl residue degradation pathway in *C. testosteroni* TA441.

## RESULTS

### Analysis of Intermediate Compounds Accumulated by Gene-Disrupted Mutants

*Comamonas testosteroni* TA441 degrades cholic acid, a major component of bile acids. In our previous study, we elucidated the steroidal ABCD ring degradation process in TA441(32, 33), which initiates after the removal of the C17 side chain. The removal of the C20-22 propionyl residue from the C17 side chain has been identified in several bacteria, including actinobacteria (*M. tuberculosis* H37Rv (2, 22–26), *Rhodococcus jostii* RHA1 (27, 28), *Thermomonospora curvata* (29), and *Mycolicibacterium neoaurum* DSM 44074 (30)) and proteobacteria (*Pseudomonas stutzeri* Chol1(31)), with highly conserved mechanisms among these species (Fig. 1A). In our previous studies analyzing intermediate compounds of cholic acid in cultures of a TA441 mutant, compounds shown in Fig. 2 were purified and identified using NMR and mass spectrometry (17). Some compounds were unstable and converted to other derivatives. For example, one intermediate converted to 7α,12α-dihydroxy-5β-cholan-4,17-dien-24-oic acid (**VII**) under acidic conditions, suggesting it was 7α,12α,17-trihydroxy-5β-cholan-4-en-24-oic acid (Fig. 1A and 2). A TA441 transposon mutant with an insertion in ORF44 accumulated 7α,12α-dihydroxy-5β-cholan-4-en-24-oic acid (**V**) when cultured with cholic acid (17). To investigate the role of genes near ORF44, we constructed gene-disrupted mutants targeting ORF35–55 (Fig. 1B) and analyzed cultures incubated with cholic acid using UPLC/MS. The analysis suggested that ORF40 to ORF44 were involved in degrading the propionyl residue.

The putative amino acid sequences of ORF40 to ORF44 showed respective similarities of 35% to the hydratase alpha subunit ChsH2 (3-oxo-4-pregnene-20-carboxyl-CoA hydratase; 3-oxo-23,24-bisnorchol-4,17(20)-dien-22-oyl-CoA hydratase), 31% to the dehydrogenase alpha subunit ChsE1 (3-oxo-4,17-pregnadiene-20-carboxyl-CoA dehydrogenase; 3-oxo-23,24-bisnorchol-4-en-22-oyl-CoA dehydrogenase), 44% to the beta subunit of the hydratase ChsH1, 63% to the retro-aldolase Ltp2 (17-hydroxy-3-oxo-4-pregnene-20-carboxyl-CoA retro-aldolase), and 44% to the beta subunit of the dehydrogenase ChsE2 of *M. tuberculosis* H37Rv (Fig. 1B). These enzymes, encoded by the *igr* operon (*chsE1* to *ltp2*, Fig. 1B), are well-studied for their roles in the propionyl residue degradation pathway (22–26). ChsE1 and ChsE2 form α_2_β_2_ heterotetramers with two active sites regulated by cholesterol (23, 24). ChsH2 comprises MaoC-like dehydratase and DUF35 domains (Fig. 1B), and ChsH2_MaoC_ and ChsH1 form a hydratase complex, while ChsH2_DUF35_ and Ltp2 form an aldolase complex, both as α_2_β_2_ heterotetramers (25, 29). In TA441, the MaoC-like domain and DUF35 domain were also found in ORF40. Consequently, the MaoC-like and DUF35 domains of ORF40 were analyzed separately.

To determine whether the proteins encoded by ORF40 to ORF44 function similarly to their counterparts in *M. tuberculosis*, we disrupted ORF40_MaoC_, ORF40_DUF35_, ORF41, ORF42, ORF43, and ORF44 using kanamycin resistance gene insertions (ORF40_MaoC_^-^, ORF40_DUF35_^-^, ORF41^-^, ORF42^-^, ORF43^-^, and ORF44^-^) (Table 1). Cultures of these mutants were analyzed using UPLC/MS after incubation with cholic acid. However, the complexity of the resulting peaks, due to the three hydroxyl groups (C3, C7, and C12) of cholic acid producing multiple derivatives (c.f. Fig. 2), necessitated the use of lithocholic acid (LCA) instead. LCA, which has only one hydroxyl group at C3 that is basically converted to a ketone group before C17 side-chain degradation. The mutants were individually incubated with LCA, and the resulting cultures were analyzed using UPLC/MS (Fig. 3). In the cultures of ORF41^-^ and ORF44^-^, 3-oxo-1,4-pregnedine-20-carboxylic acid (**IV’**) accumulated as the major intermediate compound. In contrast, 3-oxo-1,4,17-pregnetriene-20-carboxylic acid (**VII’**) was detected in the cultures of ORF40_MaoC_^-^, ORF40_DUF35_^-^, ORF42^-^, and ORF43^-^, with **VII’** being the predominant compound in the cultures of ORF40_MaoC_^-^ and ORF42^-^.

**Fig. 3.**
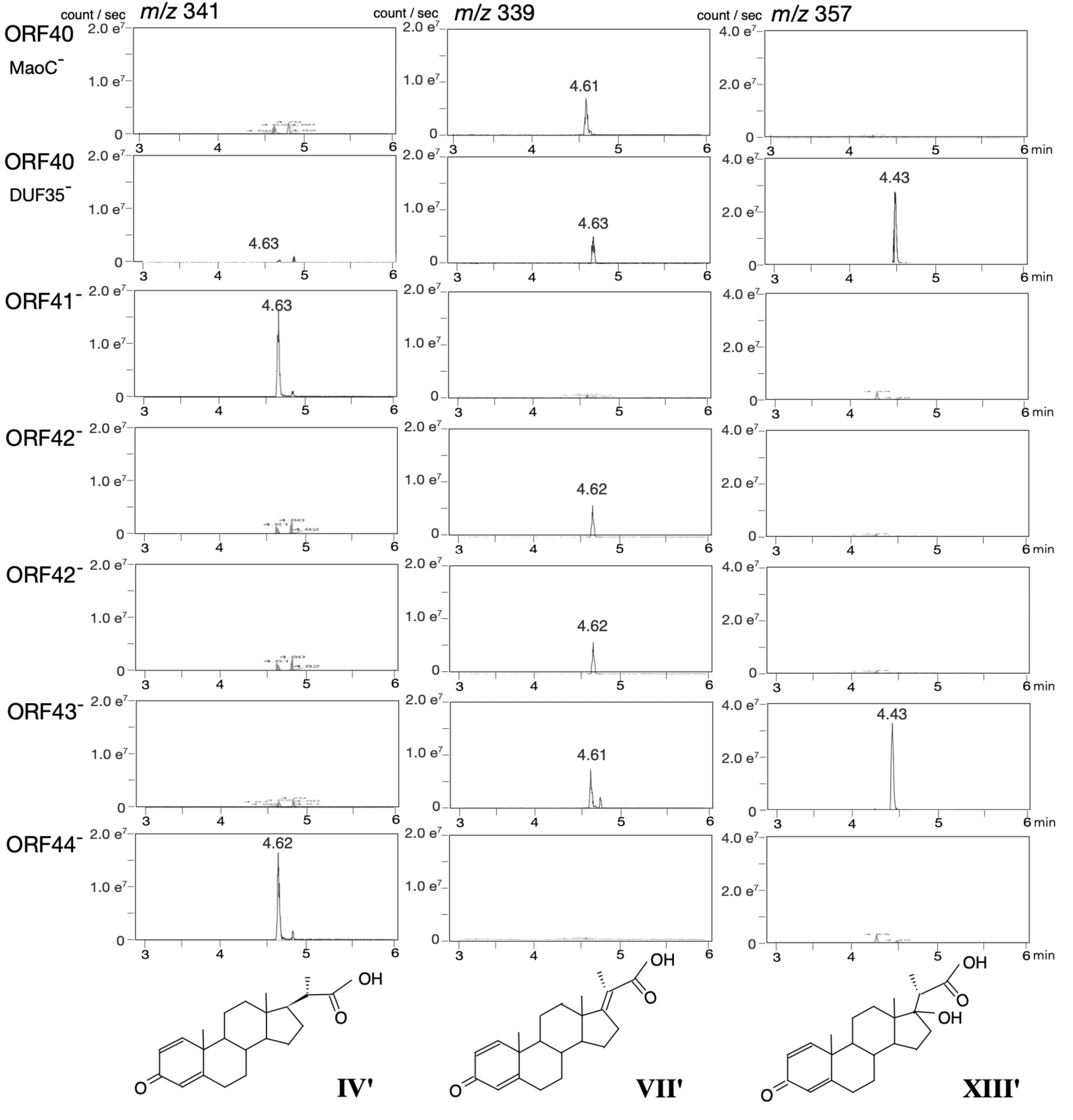
Analysis of TA441 mutants disrupted in ORF40 to ORF44, incubated with 0.05% lithocholic acid for 7 d. Mutants were constructed by inserting a kanamycin resistance gene. For ORF40, separate mutants targeting the MaoC domain (ORF40_MaoC_^−^) and DUF35 domain (ORF40_DUF35_^−^) were created. Peaks in mass spectra represent: *m/z* 341 at RT = 4.6 min (3-oxo-4-pregnene-20-carboxylic acid, **IV’**); *m/z* 339 at RT = 4.6 min (3-oxo-4,17-pregnadiene-20-carboxylic acid, **VII’**); and *m/z* 357 at RT = 4.4 min (17-hydroxy-3-oxo-1,4-pregnatriene-20-carboxylic acid, **XIII’**). The vertical axis indicates intensity (counts/sec), and the horizontal axis indicates retention time (min).

**Table 1.** strains.

| strains | Characteristics | Source or reference |
| --- | --- | --- |
| TA441 | Wild-type | (36) |
| ORF40<sub>MaoC</sub><sup>-</sup> | ORF40: :Km<sup>r</sup> mutant (inserted in MaoC domain) of TA441 | this work |
| ORF40<sub>DUF35</sub><sup>-</sup> | ORF40: :Km<sup>r</sup> mutant (inserted in DUF35 domain) of TA441 | this work |
| ORF41<sup>-</sup> | ORF41: :Km<sup>r</sup> mutant of TA441 | this work |
| ORF42<sup>-</sup> | ORF42: :Km<sup>r</sup> mutant of TA441 | this work |
| ORF43<sup>-</sup> | ORF43: :Km<sup>r</sup> mutant of TA441 | this work |
| ORF44<sup>-</sup> | ORF44: :Km<sup>r</sup> mutant (inserted in *EcoRV* site) of TA441 | this work |
| ORF40<sup>-</sup>-43<sup>-</sup> | ORF40, 41, 42, 43: :Km<sup>r</sup> mutant of TA441 | this work |

Km<sup>r</sup> : Km-resistance, (36) Arai et al. 1998. Microbiology **144**: 2895-2903

Additionally, 17-hydroxy-3-oxo-1,4-pregnedine-20-carboxylic acid (**XIII’**) accumulated in large amounts in the cultures of ORF40_DUF35_^-^ and ORF43^-^. **VII’** was detected in the cultures of ORF40_DUF35_^-^ and ORF43^-^, likely because the excessive accumulation of **XIII’** hindered the conversion of **VII’** to **XIII’**. The detected amount of **VII’** was relatively small compared to **IV’** and **XIII’**, consistent with our previous study in which **VII** was identified as an unstable intermediate (17). During isolation, **VII** rapidly degraded, making its identification challenging. This instability could partially explain the smaller amounts of **VII’** observed in this study. In TA441, the major intermediate compounds were derivatives with two double bonds at C1 and C4 in the A-ring. This contrasts with *M. tuberculosis*, where intermediates with a single double bond at C4 are predominant.

These findings support the hypothesis derived from homology searches that ORF41 and ORF44 encode 3-oxo-4-pregnene-20-carboxyl-CoA dehydrogenase (ChsE1-E2 in *M. tuberculosis*), ORF42 and ORF40_MaoC_ encode 3-oxo-4,17-pregnadiene-20-carboxyl-CoA hydratase (ChsH1-H2_MaoC_), and ORF43 and ORF40_DUF35_ encode an aldolase responsible for removing the isopropyl-CoA side chain (Ltp2-ChsH2_DUF35_). ORF40_DUF35_ is indispensable for aldolase activity, while the hydratase encoded by ORF42 and ORF40_MaoC_ remains active even in the absence of the DUF35 domain of ORF40. To better understand the specific roles and interactions of these subunits, we conducted complementation experiments to further explore their functions and interdependencies.

### Complementation Experiments Involving ORF40_MaoC_^−^ to ORF44^−^

To investigate the roles of ORF40 to ORF44, complementation experiments were conducted using mutants carrying plasmids derived from the broad-host-range vector pMFYMhpRA (33), a derivative of pMFY42 (34). The mutants were constructed as follows: ORF40_MaoC_^−^ carrying pMFYMhpRA (negative control), ORF40_MaoC_^−^ carrying a plasmid encoding ORF40_MaoC_ (pMFYMhpORF40_MaoC_), ORF40_DUF35_^−^ carrying pMFYMhpRA (negative control), ORF40_DUF35_^−^ carrying pMFYMhpORF40_DUF35_, and similarly for ORF41^−^, ORF42^−^, ORF43^−^, and ORF44^−^ with their respective complementing plasmids (Table 2 and Fig. 4. Peaks were undetectable when the data was not presented in Fig. 4).

**Fig. 4.**
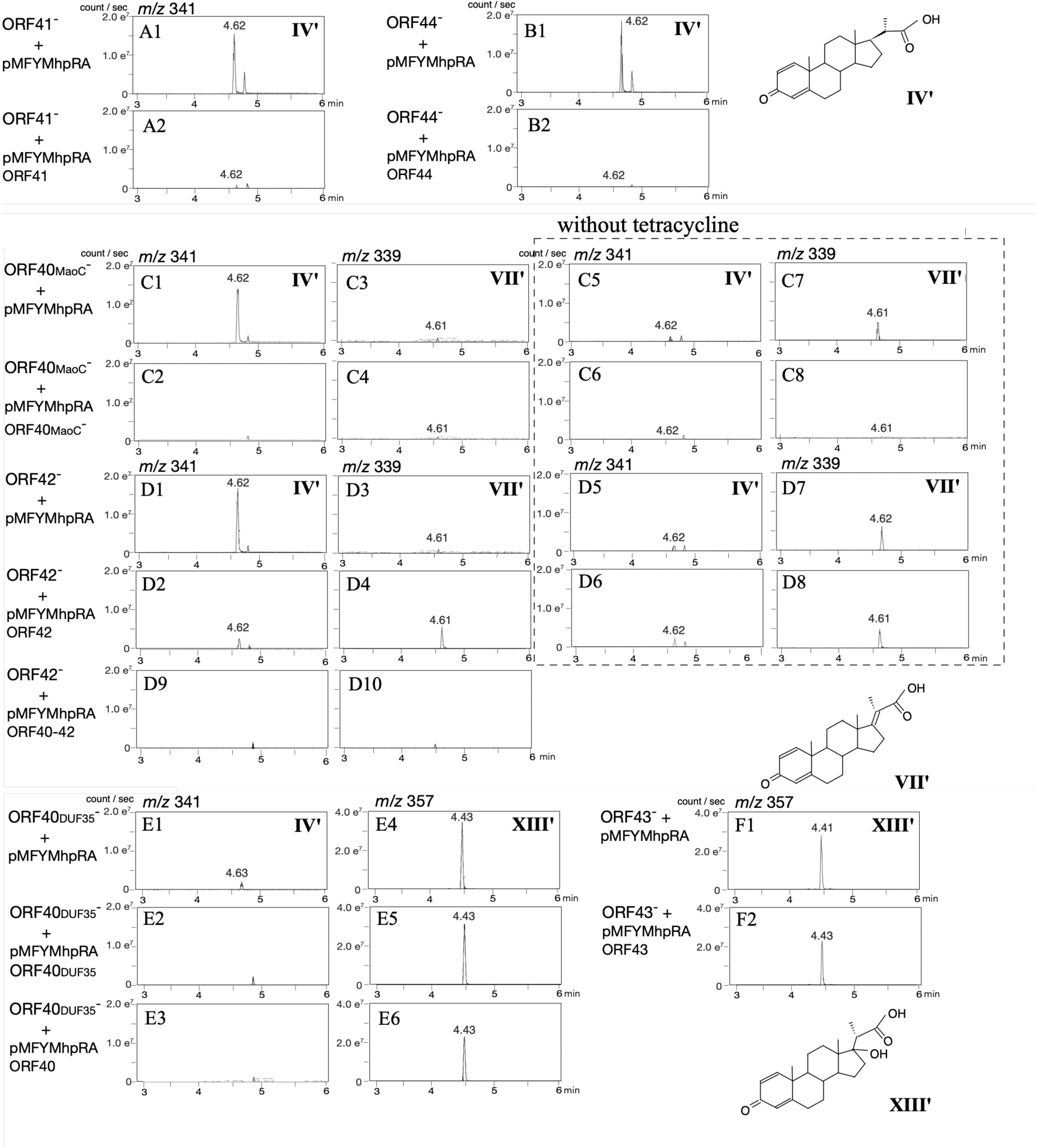
Complementation experiments with mutants disrupted in ORF40 to ORF44, incubated with 0.05% lithocholic acid for 7 d. Mass chromatograms of each mutant are presented, showing peaks at *m/z* 341 (**IV’**), *m/z* 339 (**VII’**), and *m/z* 357 **(XIII’**). Peaks were undetectable when the data was not presented. The chromatograms within broken squares represent cultures without tetracycline. Mutants are; ORF41^-^ carrying pMFYMhpRA (ORF41^-^ with pMFYMhpRA) (A1) and pMFYMhpORF41 (pMFYMhpRA-derivative carrying ORF41) (A2), ORF44^-^ carrying pMFYMhpRA (ORF44^-^ with pMFYMhpRA) (B1) and pMFYMhpORF44 (pMFYMhpRA-derivative carrying ORF44) (B2), ORF40_MaoC_^-^ carrying pMFYMhpRA (ORF40_MaoC_^-^ with pMFYMhpRA) (C1, C3, C5, and C7), ORF40_MaoC_^-^ carrying pMFYMhpORF40 (pMFYMhpRA-derivative with ORF40_MaoC_) (C2, C4, C6, and C8), ORF42^-^ carrying pMFYMhpRA (ORF42^-^ with pMFYMhpRA) (D1, D3, D5, and D7), ORF42^-^ carrying pMFYMhpORF42 (pMFYMhpRA-derivative with ORF42) (D2, D4, D6, and D8), ORF42^-^ carrying pMFYMhpORF40-42 (pMFYMhpRA-derivative with ORF40-42) (D9 and D10), ORF40_DUF35_^-^ carrying pMFYMhpRA (ORF40_DUF35_^-^ with pMFYMhpRA) (E1 and E4), ORF40 _DUF35_^-^ carrying pMFYMhpORF40 _DUF35_ (pMFYMhpRA-derivative with ORF40_DUF35_) (E2 and E5), ORF40 _DUF35_^-^ carrying pMFYMhpORF40_DUF35_ (pMFYMhpRA-derivative with ORF40) (E3 and E6), ORF43 carrying pMFYMhpRA (ORF43^-^ with pMFYMhpRA) (F1), and ORF43^-^ carrying pMFYMhpORF43 (pMFYMhpRA-derivative with ORF43) (F2).

**Table 2.** plasmids.

| plasmids | Characteristics | Source or reference |
| --- | --- | --- |
| pUC19 | Ap<sup>r</sup>, *lacZ* | (37) |
| pMFY42 | Tc<sup>r</sup>, Km<sup>r</sup>, RSF1010-based broad host range plasmid | (34) |
| pMFYMhpRA | pMFY42 derivative carrying *mhpR* and the promoter of *mhp* genes | (33) |
| pMFYMhpORF40 | pMFYMhpRA derivative carrying ORF40 | this work |
| pMFYMhpORF41 | pMFYMhpRA derivative carrying ORF41 | this work |
| pMFYMhpORF42 | pMFYMhpRA derivative carrying ORF42 | this work |
| pMFYMhpORF43 | pMFYMhpRA derivative carrying ORF43 | this work |
| pMFYMhpORF44 | pMFYMhpRA derivative carrying ORF44 | this work |
| pMFYMhpORF4041 | pMFYMhpRA derivative carrying ORF40 and 41 | this work |
| pMFYMhpORF4142 | pMFYMhpRA derivative carrying ORF41 and 42 | this work |
| pMFYMhpORF40-42 | pMFYMhpRA derivative carrying ORF40 to 42 | this work |
| pMFYMhpORF40-43 | pMFYMhpRA derivative carrying ORF40 to 43 | this work |
| pMFYMhpORF40<sub>MaoC</sub> | pMFYMhpRA derivative carrying MaoC domain of ORF40 | this work |
| pMFYMhpORF40<sub>DUF35</sub> | pMFYMhpRA derivative carrying DUF35 domain of ORF40 | this work |
| pHSG10-2-8-43 | pHSG397 derivative carrying *EcoRI* fragment containing ORF35 to 39 | this work |
| pHSG10-2-8-28 | pHSG397 derivative carrying *EcoRI* fragment containing ORF41 to 45 | this work |
| pUC37-42 | pUC derivative carrying DNA fragment containing ORF37 to 42 | this work |
| pUCORF40-Km<sup>r</sup> | pUC19 derivative carrying DNA fragment containing ORF40<sub>MaoC</sub>: :Km<sup>r</sup> | this work |
| pUCORF41-Km<sup>r</sup> | pUC19 derivative carrying DNA fragment containing ORF41: :Km<sup>r</sup> | this work |
| pUCORF42-Km<sup>r</sup> | pUC19 derivative carrying DNA fragment containing ORF42: :Km<sup>r</sup> | this work |
| pUCORF43-Km<sup>r</sup> | pUC19 derivative carrying DNA fragment containing ORF43: :Km<sup>r</sup> | this work |
| pUCORF44-Km<sup>r</sup> | pUC19 derivative carrying DNA fragment containing ORF44: :Km<sup>r</sup> (*EcoRV* site) | this work |
| pUCORF40<sub>DUF35</sub>-Km<sup>r</sup> | pUC19 derivative carrying DNA fragment containing ORF40<sub>DUF35</sub>: :Km<sup>r</sup> | this work |
| pUCORF40-43-Km<sup>r</sup> | pUC19 derivative carrying DNA fragment containing ORF40-43: :Km<sup>r</sup> | this work |

Km<sup>r</sup> : Km-resistance pSuperCosI\* (Stratagene, CA) (37) Vieira, J., and Messing, J. 1987. Methods Enzymol. **153**: 3-11. (34) Nagata Y. et al. 1993. J Bacteriol **175**:6403-6410 (33) Horinouchi M, et al. 2023. Appl Environ Microbiol **89**:e0105023.

In the cultures of ORF41^−^ and ORF44^−^ mutants carrying pMFYMhpRA (negative controls), **IV’** accumulated as the major intermediate (Fig. 4A1, 4B1), and **IV’** was degraded in the complemented mutants carrying pMFYMhpORF41 and pMFYMhpORF44 (Fig. 4A2, 4B2). Similarly, **XIII’** accumulated in ORF40_DUF35_^−^ and ORF43^−^ mutants carrying pMFYMhpRA (Fig. 4E4, 4F1). The amount of **XIII’** was reduced in cultures of ORF40_DUF35_^−^ carrying pMFYMhpORF40 and ORF43^−^ carrying pMFYMhpORF43, although the reduction was modest (Fig. 4E5, 4F2). To further investigate, we constructed a new mutant, ORF40_DUF35_^−^ carrying pMFYMhpORF40, and observed a bit more pronounced reduction in **XIII’** compared to pMFYMhpORF40_DUF35_ (Fig. 4E6). To our surprise, **IV’** unexpectedly accumulated in the cultures of the ORF40_MaoC_^-^ carrying pMFYMhpRA and the ORF42^-^ carrying pMFYMhpRA (Fig. 4C1 and 4D1), even though **VII’** had been observed as the major accumulated compound in previous experiments with these mutants. This discrepancy was thought to be attributed to the presence of tetracycline, which was added to maintain the plasmids in this experiment. To confirm this, the same experiments were repeated without tetracycline (Fig. 4C5 to C8 and 4D5 to D8). Under these conditions, **VII’** was the major compound detected in the cultures of ORF40_MaoC_^-^ carrying pMFYMhpRA and the ORF42^-^ carrying pMFYMhpRA (Fig. 4C7 and 4D7).

Tetracycline functions by inhibiting protein synthesis in bacteria through its interaction with the bacterial 30S ribosomal subunit, while the tetracycline-resistance gene encodes a pump that expels tetracycline from the cell. If tetracycline had inhibited the expression of the dehydrogenase encoded by ORF41 and ORF44, **IV’** should have accumulated in the ORF40_DUF35_^-^ and ORF43^-^ carrying pMFYMhpRA. Therefore, possibility of direct inhibition of the dehydrogenase by tetracycline was denied.

The mechanism underlying the accumulation of **IV’** in the cultures of ORF40_MaoC_^-^ and ORF42^-^ carrying pMFYMhpRA remains unclear. However, it is possible that interactions between the dehydrogenase encoded by ORF41 and ORF44 and the hydratase encoded by ORF40_MaoC_ and ORF42 influence this outcome. In the culture of ORF40_MaoC_^-^ complemented with pMFYMhpORF40_MaoC_, with tetracycline, as well as without tetracycline, the levels of both **VII’** and **IV’** were reduced to nearly undetectable levels (Fig. 4C2, 4C4, and 4C8). Similarly, in the cultures of the ORF42^-^ carrying pMFYMhpRA with tetracycline, **IV’** levels were barely detectable after complementation with pMFYMhpORF42 (Fig. 4D2). However, the reduction in **VII’** levels was minor in the cultures of ORF42^-^ with pMFYMhpORF42 (Fig. 4D4 and 4D8). Considering the possibility that ORF40_MaoC_- and ORF42-encoded enzymes did not properly form the complex, the ORF42^-^ was complemented with a plasmid encoding ORF40 to ORF42 (pMFYMhpORF40-42), and under these conditions, both **IV’** and **VII’** were nearly undetectable (Fig. 4D9 and 4D10).

While the precise reasons for these observations on ORF40_MaoC_^-^ and ORF42^-^ remain unclear, the complementation experiments support the hypothesis that the dehydrogenase encoded by ORF41 and ORF44 converts **IV’**-CoA to **VII’**-CoA. This is followed by the addition of a water molecule at C17, catalyzed by the hydratase encoded by ORF42 and ORF40_MaoC_, and the removal of the C17 residue as a ketone group, mediated by the aldolase encoded by ORF43 and ORF40_DUF35_. These results also suggest the possibility of interactions between the enzyme complexes, which may play a crucial role in the sequential degradation of the C17 side chain.

### Complementation Experiments with ORF40–43 Disrupted Mutant

For further analysis, we attempted to construct a mutant with a complete disruption of ORF40–44. Despite multiple attempts using various plasmids designed for homologous recombination, all efforts to disrupt the entire ORF40–44 region were unsuccessful. Consequently, we constructed an ORF40–43 disrupted mutant (ORF40^−^–43^−^) as an alternative. This mutant serves as a suitable replacement for ORF40–44 disrupted mutants to investigate whether all three enzymes are indispensable for the removal of the propionyl residue, as the absence of either ORF41 or ORF44 abolishes dehydrogenase activity (Fig. 3 and 4). Then we respectively introduced pMFYMhpRAORF41, a pMFYMhpRA-based plasmid encoding ORF40 and 41 (pMFYMhpORF4041), a pMFYMhpRA-based plasmid encoding ORF41 and 42 (pMFYMhpORF4142), pMFYMhpORF40-42, a pMFYMhpRA-based plasmid encoding ORF40 to 43 (pMFYMhpORF40-43), and pMFYMhpRA (negative control) into the ORF40^−^-43^−^ (table 1 and 2). The resulting mutants were incubated with LCA in the presence of tetracycline (Fig. 5). As observed in previous experiments, mutants lacking either ORF40 or ORF42 primarily accumulated **IV’**, except for ORF40^−^–43^−^ carrying pMFYMhpORF4041. This mutant, expressing ORF40, ORF41, and ORF44, accumulated a small amount of **VII’**, indicating partial dehydrogenase activity even in the presence of tetracycline. Independent colonies of this mutant consistently produced similar results, confirming its partial activity in converting **IV’** to subsequent intermediates.

**Fig. 5.**
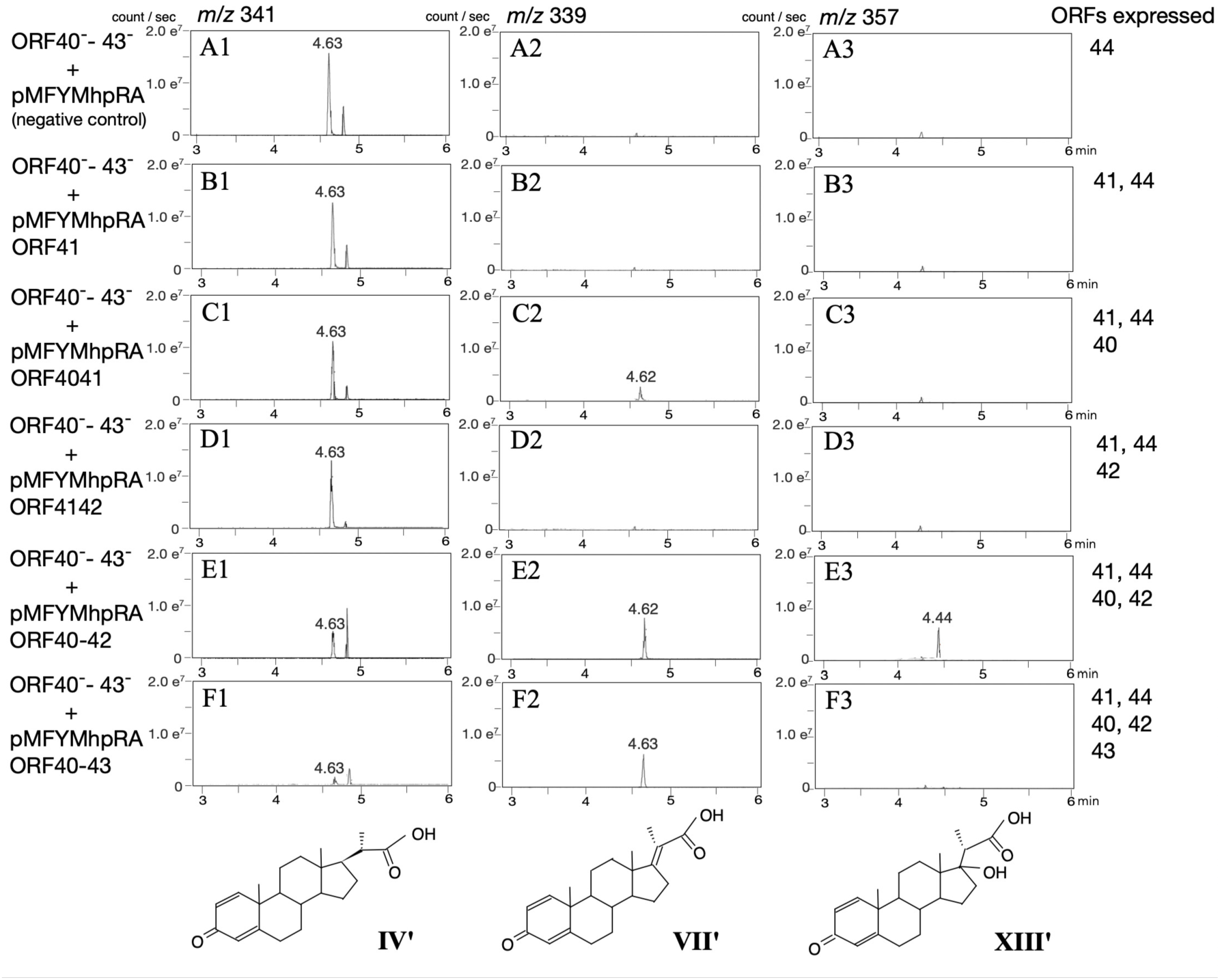
Complementation experiments with ORF40–43 disrupted mutants (ORF40^-^-43^-^) incubated with 0.05% lithocholic acid for 1 d, followed by the addition of 3-(3-hydroxyphenyl)propionic acid (3HPP) for gene induction, and further incubation for 6 d. Mass spectra display peaks at *m/z* 341 (**IV’**), *m/z* 339 (**VII’**), and *m/z* 357 (**XIII’**). ORFs expressed in each mutant are indicated on the right. Mutants are; ORF40^-^-43^-^ carrying pMFYMhpRA (A), carrying pMFYMhpORF41 (B), carrying pMFYMhpORF4041 (pMFYMhpRA-derivative with ORF40 and 41) (C), carrying pMFYMhpORF4142 (pMFYMhpRA-derivative with ORF41 and 42) (D), carrying pMFYMhpORF40-42 (pMFYMhpRA-derivative with ORF40, 41, and 42) (E), and carrying pMFYMhpORF40-43 (pMFYMhpRA-derivative with ORF40 to 43) (F).

In the culture of ORF40^−^–43^−^ carrying pMFYMhpORF40–42, which expresses ORF40, ORF41, ORF42, and ORF44, **IV’** was converted to **VII’** and **XIII’** (Fig. 5F), and cultures of ORF40^−^–43^−^ carrying pMFYMhpORF40–43, expressing all ORF40 to ORF43 proteins, showed nearly undetectable levels of **IV’** and **XIII’**, though **VII’** remained detectable (Fig. 5H). This result is consistent with previous findings using ORF42^−^ carrying pMFYMhpORF42 (Fig. 4D4 and D8), where **VII’** conversion was incomplete. These findings suggest that while the enzyme complex encoded by ORF40– 43 is capable of catalyzing the reactions, some inefficiencies remain, potentially due to limitations in substrate processing or enzyme interactions.

## DISCUSSION

In this study, we identified the functions of ORF40–44 encoded proteins in *C. testosteroni* TA441 by constructing gene-disrupted mutants and comparing them with the corresponding Chs enzymes in *M. tuberculosis*. ORF41 and ORF44 were shown to encode the dehydrogenase responsible for converting 3-oxo-1,4-pregnediene-20-carboxyl-CoA to 3-oxo-1,4,17-pregnatriene-20-carboxyl-CoA (ChsE1-ChsE2 in *M. tuberculosis*). ORF42 and ORF40_MaoC_ encode the hydratase that adds a water molecule at C17 (ChsH1-ChsH2_MaoC_), while ORF43 and ORF40_DUF35_ encode the aldolase that removes the isopropyl-CoA side chain at C17 (Ltp2-ChsH2_DUF35_).

Proteobacteria such as TA441 and *P. stutzeri* Chol1 primarily produce intermediates with two double bonds at C1 and C4 and a ketone group at C3 during steroid degradation. In contrast, Actinobacteria, including *M. tuberculosis*, typically produce intermediates with a double bond at C4 and a ketone group at C3. This difference likely arises because Actinobacteria can introduce a hydroxyl group at C9 of intermediates before removing the C17 side chain and when they are with two double bonds at C1 and C4 and a ketone group at C3, leading to automatic A-ring aromatization and B-ring cleavage.

ORF40_DUF35_ was found indispensable for aldolase activity, while ORF42-40_MaoC_ hydratase retained activity without the DUF35 domain of ORF40. Despite the low amino acid identity (30–45% except for Ltp2) between TA441 and *M. tuberculosis* enzymes, their functional and structural similarities are notable. Based on these findings, we propose naming ORF40–44 of TA441 as *chsH2*, *chsE1*, *chsH1*, *ltp2*, and *chsE2*, respectively.

The three-dimensional (3D) structures of (ChsE1-ChsE2)_2_ dehydrogenase, (ChsH1-ChsH2_MaoC_)_2_ hydratase, and (Ltp2-ChsH2_DUF35_)_2_ aldolase of *M. tuberculosis* have been elucidated through crystal structure analyses (25, 26, 29). Using AlphaFold (35), we predicted the 3D structure of the (ChsH1-ChsH2_MaoC_)_2_ and (Ltp2-ChsH2_DUF35_)_2_ complexes in *C. testosteroni* TA441, as shown in Fig. 6A1 (with expected position errors illustrated in supplementary material Fig. S2). This model closely resembles the docking model of the (ChsH1-ChsH2_MaoC_)_2_ complex of *M. tuberculosis* and the (Ltp2-ChsH2_DUF35_)_2_ complex of *T. curvata* (29), despite slight differences in spatial arrangement. For instance, the upper region of the (ChsH1-ChsH2_MaoC_)_2_ complex in TA441 is slightly wider than that of *M. tuberculosis*, and the spacing between the (ChsH1-ChsH2_MaoC_)_2_ and (Ltp2-ChsH2_DUF35_)_2_ complexes is also greater. Comparison with AlphaFold models of *M. tuberculosis*’s (ChsH1-ChsH2-Ltp2)_2_ complex revealed that the (Ltp2-ChsH2_DUF35_)_2_ complex in *M. tuberculosis* is rotated counterclockwise by ∼90° relative to the (ChsH1-ChsH2_MaoC_)_2_ complex (Fig. 6A1 and 6B1). This angular difference will not impact catalytic activity, as experiments showed that separately expressed ChsH2_MaoC_ and ChsH2_DUF35_ retained hydratase and aldolase activities (predicted structure; Fig. 6A4 and experimental data; Fig. 4C4). Another 3D-model of (ChsH1-ChsH2-Ltp2) complex of *M. tuberculosis* was presented based on small-angle X-ray scattering (SAXS) and single particle electron microscopy data (26). In this model, ChsH1-ChsH2 and Ltp2 of *M. tuberculosis* form a protomer and two protomers were docked with Rosetta local refine to present a model in which two protomers joined symmetry with respect to a point at Ltp2. However, all the five AlphaFold presented models of (ChsH1-ChsH2-Ltp2) _2_ complex of TA441 aligned with the one presented in Fig. 6A1.

**Fig. 6.**
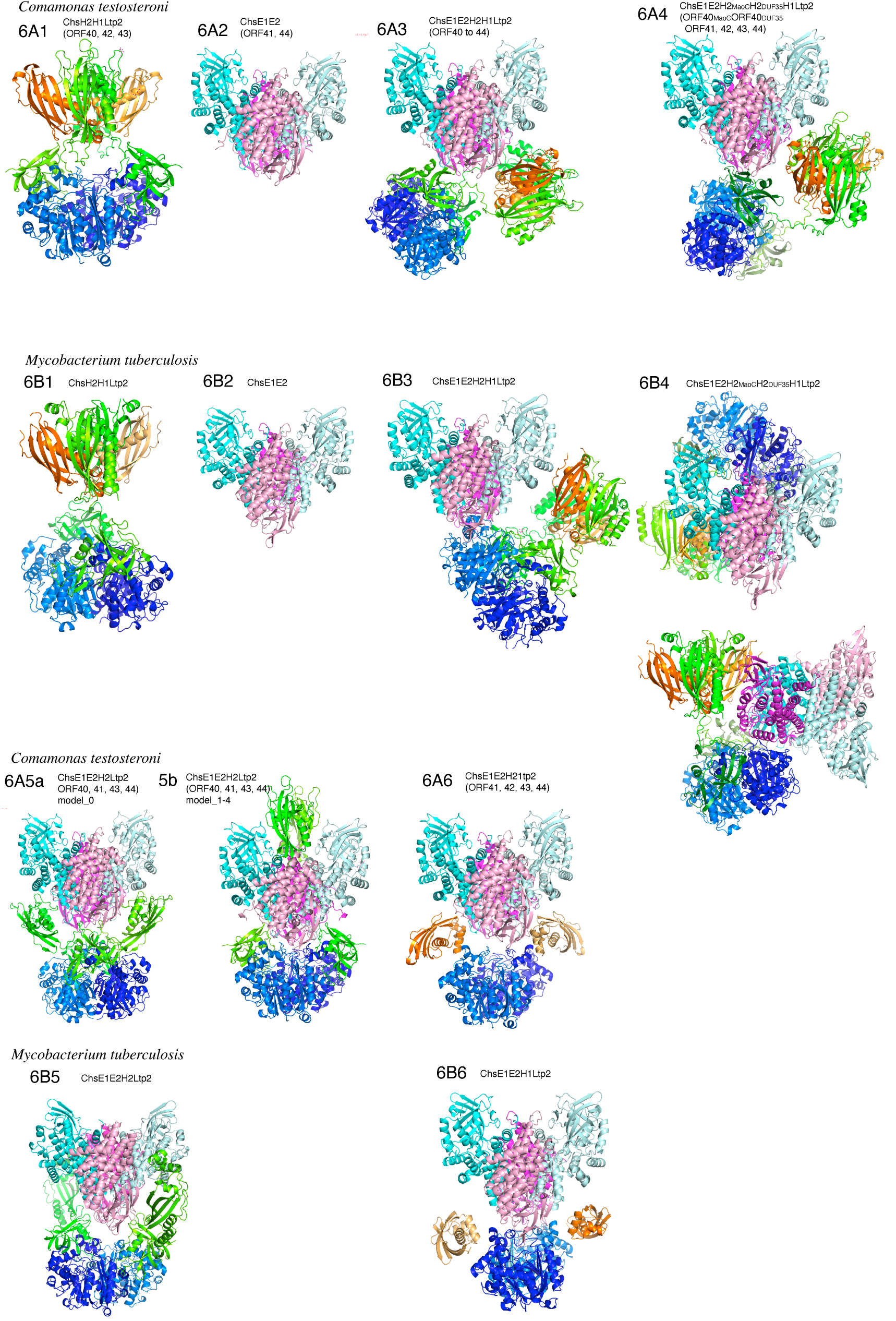
AlphaFold-predicted structures of ORF40 to 44 encoded proteins in *C. testosteroni* TA441 (6A) and corresponding enzymes in *M. tuberculosis* (6B). Complexes modeled include (ChsH1-ChsH2_MaoC_)_2_, (Ltp2-ChsH2DUF_35_)_2_, and (ChsE1-ChsE2)_2_, as well as their combined interactions. Putative structures of proteins in other mutants are shown in supplementary materials Fig. S1. Expected positional errors are shown in supplementary Fig. S2. Color scheme: cyan/light cyan: ChsE1, magenta/light magenta: ChsE2, orange/light orange: ChsH1, green/light green: ChsH2 (or ChsH2_MaoC_ when divided),, moss green/light moss green: ChsH2_DUF_, and blue/light blue: Ltp2. Figures A1B1; (ChsH1_2_-ChsH2_MaoC_)_2_-(Ltp2-ChsH2_DUF35_)_2_, A2B2; (ChsE1-ChsE2)_2_, A3B3; (ChsE1-ChsE2)_2_ with (ChsH1_2_-ChsH2_MaoC_)_2_-(Ltp2-ChsH2_DUF35_)_2_, A4B4; (ChsE1-ChsE2)_2_ with (ChsH1_2_-ChsH2_MaoC_)_2_-(Ltp2-ChsH2_DUF35_)_2_, ChsH2_MaoC_ and ChsH2_DUF35_ are separately expressed A5B5; (ChsE1-ChsE2)_2_ with ChsH2_2_ and Ltp2_2_, and A6B6; (ChsE1-ChsE2)_2_with ChsH1_2_ and Ltp2_2_. 6B4 shows the same model with different angle. 6A5a is the first model and 6A5b is the second model of (ChsE1-ChsE2)_2_with ChsH1_2_and Ltp2_2_of TA441 presented by Alphafold. The third to fifth were similar to the second one.

AlphaFold model of the (ChsE1-ChsE2)_2_ dehydrogenase of TA441 (Fig. 6A2) was also found to be structurally similar to its counterpart in *M. tuberculosis* (Fig. 6B2). Our experimental data obtained in this study suggested some interaction between (ChsH1-ChsH2-Ltp2)_2_ and (ChsE1-ChsE2)_2_ complexes (Fig. 4C and 4D). Therefore, we performed Alphafold modeling with all the subunits, ChsH1, ChsH2, Ltp2, ChsE1 and ChsE2 (Fig. 6A3). When all the subunits were analyzed together, AlphaFold modeling revealed that the (ChsE1-ChsE2)_2_ dehydrogenase complex is positioned so close to Ltp2 that it causes a distortion in the (ChsH1-ChsH2_MaoC_)_2_-(Ltp2-ChsH2_DUF35_)_2_ complex (Fig. 6A3). This suggests that, although (ChsE1-ChsE2)_2_ does not form a stable complex with (ChsH1-ChsH2_MaoC_)_2_-(Ltp2-ChsH2_DUF35_)_2_, some degree of interaction occurs. Such an interaction may be necessary for efficient removal of the propionyl residue at C17.

When ChsH2_MaoC_ and ChsH2_DUF35_ were expressed separately, the distortion in the (ChsH1-ChsH2_MaoC_)_2_-(Ltp2-ChsH2_DUF35_)_2_ complex became even more pronounced (Fig. 6A4). Despite this structural alteration, the hydratase and aldolase activities were maintained, as observed in Fig. 4C1 and 4C2. In contrast, when ChsH1 or ChsH2_MaoC_ were disrupted, preventing the formation of the (ChsH1-ChsH2_MaoC_)_2_-(Ltp2-ChsH2_DUF35_)_2_ complex, the AlphaFold models indicated that (ChsE1-ChsE2)_2_ moved closer to Ltp2, occupying the position where (ChsH1-ChsH2_MaoC_)_2_ would normally be located (Fig. 6A5ab and 6A6). This configuration further supports the hypothesis that (ChsE1-ChsE2)_2_ interacts with Ltp2 and that these interactions may play a role in catalysis. Notably, experimental results demonstrated that the dehydrogenase activity of (ChsE1-ChsE2)_2_ was abolished under tetracycline treatment when ChsH1 or ChsH2_MaoC_ were disrupted (Fig. 4C1 and 4D1). This suggests that the proper positioning of (ChsH1-ChsH2_MaoC_)_2_ may prevent (ChsE1-ChsE2)_2_ from approaching Ltp2 inappropriately, thereby ensuring optimal enzymatic activity. The fact that ChsH2_MaoC_ and ChsH2_DUF35_ are connected as a single protein may stabilize the interaction network and regulate the spatial positioning of the three complexes. AlphaFold models of *M. tuberculosis* revealed a similar phenomenon. When ChsH2_MaoC_ and ChsH2_DUF35_ were modeled separately, (ChsE1-ChsE2)_2_ was positioned to produce distortion in the (ChsH1-ChsH2_MaoC_)_2_-(Ltp2-ChsH2_DUF35_)_2_ complex (Fig. 6B4). ChsH1 or ChsH2_MaoC_ disruption also caused improper positioning of (ChsE1-ChsE2)_2_ and Ltp2_2_ (Fig. 6B5 and 6B6) as it did in TA441.

The structural and functional similarities between TA441 and *M. tuberculosis* enzymes are particularly striking given the relatively low amino acid sequence identities (30– 45% for most subunits, except Ltp2 at ∼60%). This highlights the evolutionary conservation of the steroid degradation pathway. We previously identified the genes and pathways involved in the degradation of steroidal A-, B-, C-, and D-rings in TA441 (32, 33). Further comparative analyses of AlphaFold structures from other steroid-degrading bacteria, combined with experimental validation, may uncover universal mechanisms governing steroid degradation and its ecological and biomedical significance.

## Abbreviations

UPLC/MS: ultra-performance liquid chromatography/mass spectrometry
RT: retention time
MW: molecular weight
CoA: coenzyme A
3HPP: 3-(3-Hydroxyphenyl)propionic acid
PCR: polymerase chain reaction

## MATERIALS AND METHODS

### Culture conditions

Mutant strains of *Comamonas testosteroni* TA441 were cultured at 30°C in a medium composed of equal volumes of Luria-Bertani (LB) medium and C medium, a mineral medium optimized for TA441 (9). This mixed medium was used because it facilitates the accumulation of intermediate compounds in mutants more effectively than either C medium or LB medium alone (unpublished data). Lithocholic acid (LCA) was added as a filter-sterilized solution in DMSO, with a final concentration of 0.05% (w/v). Similarly, 3-(3-Hydroxyphenyl)propionic acid (3HPP) was prepared as an acetonitrile solution and added to the medium at a final concentration of 0.1% (w/v). Construction of Deletion Mutants, Plasmids, and Mutants for Complementation Experiments

To construct the ORF40_MaoC_– mutant (Table 1), an *Hpa*I site was introduced into ORF40_MaoC_ on plasmid pHSG10-2-8-28 (Table 2) using primers ORF40_Km^r^ and ORF40_Km^r^RC (Table 3). The kanamycin resistance gene (Km^r^) was subsequently inserted into the *Hpa*I site, resulting in the construction of plasmid pUCORF40-Km^r^ (ORF40::Km^r^). The plasmid was introduced into *C. testosteroni* TA441 cells via electroporation, and successful transformants were selected on LB plates containing kanamycin. The insertion of Km^r^ into ORF40 was confirmed by PCR amplification using DNA extracted from the resultant transformants.

**Table 3.**
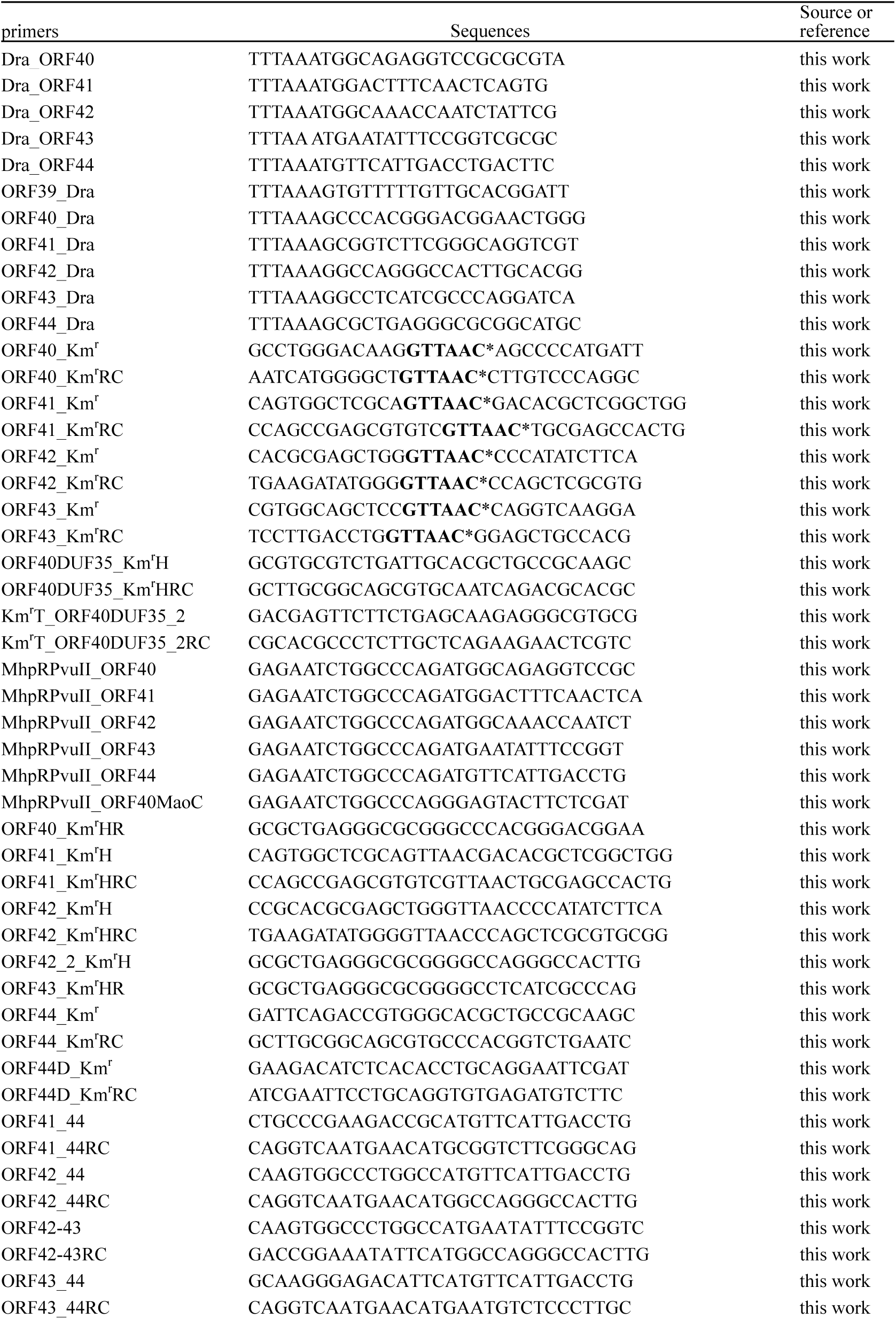
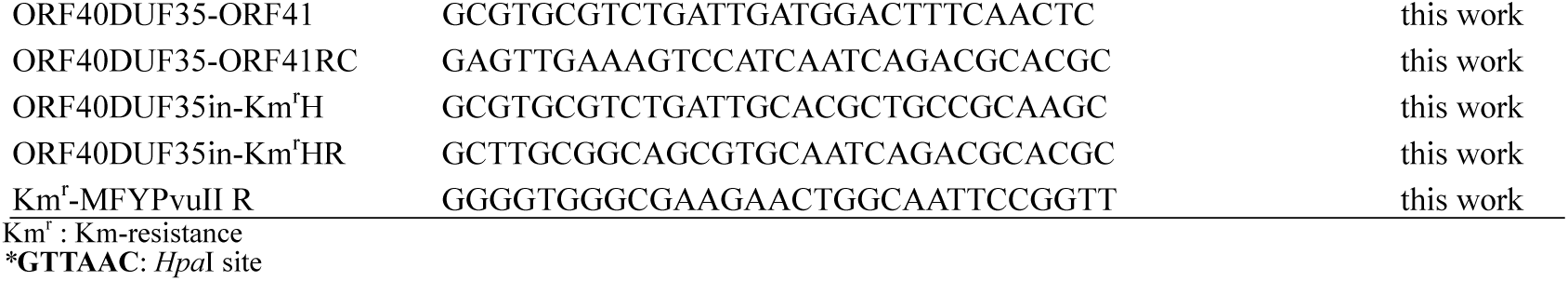
primers.

Deletion mutants for ORF41 through ORF43 were constructed using similar methods with appropriate plasmids and primers listed in Tables 2 and 3. For the construction of pUCORF44-Km^r^, the Km^r^ gene was inserted into the *Eco*RV site within ORF44. To disrupt ORF40 through ORF43, plasmid pUCORF40-43-Km^r^ was constructed by connecting a DNA fragment PCR-amplified from pUC37-42 using primers ORF39_Dra and Dra_ORF42, with a DNA fragment PCR-amplified from pUCORF43-Km^r^ using primers Dra_ORF43 and ORF44_Dra.

For pUCORF40_DUF35_-Km^r^, two DNA fragments were amplified: one from the Km^r^ gene using primers ORF40_DUF35__Km^r^H and Km^r^T_ORF40_DUF35__2RC, and another from pUCORF37-42 using primers Km^r^T_ORF40_DUF35__2 and ORF40_DUF35__Km^r^HRC. These fragments were assembled using the In-Fusion HD Cloning Kit (TAKARA, Japan).

Plasmids for complementation experiments, such as pMFY40 and pMFY40_DUF35_ (Table 2), were constructed by amplifying DNA fragments containing the target ORF(s) and the Km^r^ gene with suitable primers listed in Table 3. For example, primers MhpRPvuII_ORF40 and ORF40_Km^r^HR were used to amplify the ORF40 fragment, while primers ORF40_Km^r^H and Km^r^-MFYPvuIIR were used to amplify the Km^r^ fragment. The amplified fragments, along with *Pvu*II-digested pMFYMhpRA, were connected using the In-Fusion HD Cloning Kit.

### Ultra-High-Performance Liquid Chromatography (UPLC)/MS

A 1 mL culture was extracted twice with an equal volume of ethyl acetate under acidic conditions (adjusted to pH 2 with HCl). The ethyl acetate layer was collected, dried, and dissolved in 1 mL of methanol. A 5 µL aliquot of the methanol solution was injected into the ultra-high-performance liquid chromatography/mass spectrometry (UPLC/MS) system, Waters Acquity UPLC H-Class-QDa (Waters, Milford, MA, USA). Chromatographic separation was performed using a reversed-phase column (BEH C18, 2.1 × 50 mm, 1.7 µm, Waters) at a flow rate of 0.6 mL/min. Elution was carried out with 10% solution A (CH_3_CN) and 90% solution B (H_2_O:HCOOH = 100:0.05) for 1.0 min, followed by a linear gradient from 10% solution A and 90% solution B to 80% solution A over 3.0 min, which was maintained for 1 min and back to 10% solution A over 1.5 min with following equilibrating for 1 min. Metabolite detection was performed using negative ion electrospray ionization.

## Data Availability

The authors affirm that materials and data reasonably requested by others will be made available from a publicly accessible collection or provided in a timely manner at reasonable cost and in limited quantities to members of the scientific community for noncommercial purposes.

## Acknowledgments

The authors sincerely thank Dr. Reizo Kato (Head of Condensed Molecular Materials Laboratory, RIKEN) and Dr. Yousoo Kim (Head of Surface and Interface Science Laboratory, RIKEN) for their thoughtful support and guidance. The authors also express their gratitude to Dr. Takemichi Nakamura and the Support Units at RIKEN (Molecular Structure Characterization Unit, RIKEN CSRS, Wako) for their invaluable assistance with UPLC/MS analysis.

## SUPPLEMENTARY MATERIAL

**Fig. S1** Additional AlphaFold models of proteins expressed in the mutants used in Figs. 4 and 5. Detected enzyme activities are shown on the right. Tc: tetracycline; *\**: not examined. Color scheme: cyan/light cyan: ChsE1, magenta/light magenta: ChsE2, orange/light orange: ChsH1, green/light green: ChsH2 (or ChsH2_MaoC_ when divided), moss green/light moss green: ChsH2_DUF_, and blue/light blue: Ltp2. S1A; ORF40_MaoC_^−^ carrying pMFYMhpRA (mutant from Fig. 4C), S1B; ORF40_DUF35_^−^ carrying pMFYMhpRA (mutant from Fig. 4E), shown from a different angle, S1C; ORF42^−^ carrying pMFYMhpRA (mutant from Fig. 4D), including another model (same as Fig. 6A5a,b), S1D; ORF40–43^−^ carrying pMFYMhpORF4041 (mutant from Fig. 5C). S1E; ORF40–43^−^ carrying pMFYMhpORF4142 (mutant from Fig. 5D), S1F; ORF40–43^−^ carrying pMFYMhpORF404142 (mutant from Fig. 5G), shown from a different angle, SG1; Models of (ChsE1-ChsE2)_2_ with (ChsH1-ChsH2_MaoC_)_2_-(Ltp2-ChsH2_DUF35_)_2_ where ChsH2_MaoC_ and ChsH2_DUF35_ are separately expressed: model_1 to model_3 (Fig. SG1, same as Fig. 6A4), model_4 (Fig. SG2), and model_5 (Fig. SG3).

**Fig. S2** Expected position error in the AlphaFold models shown in Fig. 6. Darker green indicates stronger predicted interactions between amino acid residues on the X- and Y-axes.

